# High-order interactions maintain or enhance structural robustness of a coffee agroecosystem network

**DOI:** 10.1101/2021.02.22.432328

**Authors:** Cecilia González González, Emilio Mora Van Cauwelaert, Denis Boyer, Ivette Perfecto, John Vandermeer, Mariana Benítez

## Abstract

The capacity of highly diverse systems to prevail has proven difficult to explain. In addition to methodological issues, the inherent complexity of ecosystems and issues like multicausality, non-linearity and context-specificity make it hard to establish general and unidirectional explanations. Nevertheless, in recent years, high order interactions have been increasingly discussed as a mechanism that benefits the functioning of highly diverse ecosystems and may add to the mechanisms that explain their persistence. Until now, this idea has been explored by means of hypothetical simulated networks. Here, we test this idea using an updated and empirically documented network for a coffee agroecosystem. We identify potentially key nodes and measure network robustness in the face of node removal with and without incorporation of high order interactions. We find that the system’s robustness is either increased or unaffected by the addition of high order interactions, in contrast with randomized counterparts with similar structural characteristics. We also propose a method for representing networks with high order interactions as ordinary graphs and a method for measuring their robustness.

**Highlights:**

- The robustness of a coffee-associated ecological network is either increased or unaffected by the incorporation of high order interactions.
- A method is proposed for representing high order interactions in ordinary networks.
- A method is proposed to measure the robustness of networks with high order interactions.
- High order interactions may promote the persistence of diverse ecosystems.

## **1.** Introduction

The link between an ecosystem’s diversity, structure and functioning has long been debated in ecology. Both empirical and theoretical studies have tried to decipher the nature of their relationship and the factors that take part in shaping it. On the one hand, the existence of different definitions for these features has contributed to the difficulty of the task, while on the other hand, an intrinsic complexity stems from the very numerous elements, processes and scales that interact to give rise to these qualities (Ives & Carpenter 2007). Early ideas on the topic focused on the notion of stability, and maintained that diversity made ecosystems stable through species limiting each others’ growth by predation or competition (Odum 1953; MacArthur 1955; Elton 1958). These notions were dramatically challenged by the work of Robert May (1972; 1973), who used linear stability analyses to show that communities modelled as random networks lose local stability as the number of species, the number of interactions, or their strength rise. These results caused commotion in the scientific community, as they seemed to contradict the very real biodiversity found around the world. Since then, two main extensions have helped reconcile theory with observation; mainly: the use of realistic community structures (Lawlor 1978; Lawlor 1980) and the complementation of linear stability analyses with other methods to assess ecosystem function from both a structural and a dynamical point of view like robustness, feasibility or structural stability (Landi et al. 2018). It is now generally recognized that diversity tends to positively correlate with some measures of ecosystem functioning, like stability, robustness or productivity. Nevertheless, this does not mean that diversity is the direct driver of these traits, rather, it should be regarded as an ‘umbrella’ indicator of many ecological mechanisms that are inherent to ecosystems and that are the actual determinants of the diversity-function relationships (McCann 2000). Such mechanisms and how they may favor the assembly and reproduction of highly diverse communities are now the focus of many studies (Chesson 2000; Levine et al. 2017).

Different mechanisms have since been proposed to enable the coexistence of species in highly diverse systems (Chesson 2000; Wright 2002; Adler et al. 2013; Levine et al. 2017). Recently, high order interactions (HOI) have been proposed as a key mechanism for the persistence of diverse communities (Bairey et al. 2016, Grilli et al. 2017). A HOI is the effect a species has *on the interaction* between any other two species. The importance of this kind of interactions has been recognized, as they are quite common: ecosystem engineering, predatory adaptive behavior, changes in foraging, facilitation, mutualisms and many so-called trait-mediated effects commonly involve HOIs (Beckerman et al. 1997; Werner & Peacor 2003; Holt & Barfield 2012; Kéfi et al. 2012; Bairey et al. 2016). Bairey et al. (2016) computationally explored the role of HOIs on the linear stability and feasibility of systems described as virtual random networks and found that HOIs could indeed attenuate or even revert a negative relationship between the number of species and stability.

While the findings of Bairey and coworkers (2016) and other recent theoretical work have greatly contributed to our understanding of the relationship between HOIs and species coexistence (Grilli et al., 2017; Singh & Baruah, 2020; Li et al., 2020), they rely on hypothetical networks whose interactions are set randomly and do not represent known ecological interactions, or on the assessment of some focal species (Mayfield & Stouffer, 2017). It thus remains unclear how HOIs may affect the function of empirically-documented networks which, arguably, capture some aspects of their structure and dynamics in a more faithful manner. There are now some well-studied ecological and few agroecological networks that could help fill this important gap (Scheffer 1997; Yoon et al. 2004; Fortuna et al. 2014; Perfecto and Vandermeer, 2015; López Martínez 2017). Agroecosystems cover around 40 % of the Earth’s surface (Foley et al. 2005), represent a substantial part of the world’s biodiversity, and have just recently begun to be analyzed from a network perspective (Bohan et al. 2013; López Martínez 2017). The insights gained from such a system-level approach hold the potential to guide our actions around major issues like autonomous pest control, disease outbreaks and biodiversity conservation in agricultural landscapes (Vandermeer et al. 2010, 2018; Ramos et al. 2018).

With this in mind, in the present study we updated and analyzed an empirically-based network for a coffee agroecosystem in southern Mexico. This biodiverse agroecosystem has been studied for about three decades and many of its species and interactions have been thoroughly described (Perfecto & Vandermeer 2015). Importantly, different HOIs have been found to play a key role in the dynamics of the main coffee pests and their natural enemies (Vandermeer et al. 2010; Perfecto et al., 2021), motivating discussions on different formalisms to integrate HOIs to ecological network analyses, which remain an underdeveloped area (Golubski et al. 2016; Battiston et al., 2020). Thus, we analyzed the coffee agroecosystem network from a structural perspective in order to investigate the effects of HOIs on the overall robustness of this system, defined as its capacity to remain connected in the face of node removal representing species loss.

To this aim, we propose a method for representing networks with high order interactions as ordinary graphs and a method for measuring their robustness which is a modification of Piraveenan et al. (2013). Our work aims to contribute to the understanding of the mechanisms underlying species coexistence in highly diverse systems, as well as to provide novel insights that can inform management practices based on the biological understanding of agroecosystems.

## 2. Methods

### 2.1 Study site

The study site is “Finca Irlanda”, a 320 ha coffee plantation situated on the highlands of El Soconusco, Chiapas (158110 N, 928200 W; 900 masl). Precipitation in the region averages 4500 mm per year and the vegetation type is seasonal tropical forest. Nevertheless, primary vegetation has been almost completely replaced by coffee plantations with different management intensities, aside from some tiny fragments of original forest kept in some farms. In Finca Irlanda, there is a portion of such original vegetation set aside for conservation, while the management of the surrounding productive area involves keeping the shade provided by native trees, which, among other practices, make it a highly biodiverse agroecosystem (Perfecto and Vandermeer, 2015).

It is convenient to detail some parts of the complex ecological web found in the study site. There are four main antagonists of coffee plants: the coffee leaf rust, *Hemileia vastatrix*, the coffee berry borer, *Hypothenemus hampei* (see Figure 3d further), the coffee leaf miner, *Leucoptera coffeella*, and the coffee green scale, *Coccus viridis* (Figure3c). The last one keeps a spatially clustered mutualistic relationship with ants of the *Azteca* genus (Figure 3e), which feed on the honeydew produced by the scales while protecting them from being eaten by a lady beetle, *Azya orbigera*. Thanks to this protection, the scale populations reach high levels within the clusters, which in turn increases their probability of being infected by the white halo fungus, *Lecanicillium lecanii*, a fungus that is also capable of infesting the coffee rust. By patrolling coffee plants where green scales feed, *Azteca* keeps other herbivores, like the berry borer beetle or the leaf miner from establishing big populations on these plants. However, all the effects that the *Azteca* ants have on the system are temporally inhibited by flies in the genus *Pseudacteon* (Family: Phoridae), who are parasitoids of the *Azteca* ants, and that cause them to retrieve to their nests, hide or dramatically reduce their movement whenever they sense a fly nearby. This inhibition of *Azteca* leaves the scales and the coffee plants unprotected for a period of time, a lapse that has been proven to be ecologically relevant and that for example, is enough for allowing *Azya orbigera* to prey on the scales or oviposit underneath them, ensuring nourishment for their future larvae (Liere & Larsen 2010; Vandermeer et al. 2010).

**Figure 3.**
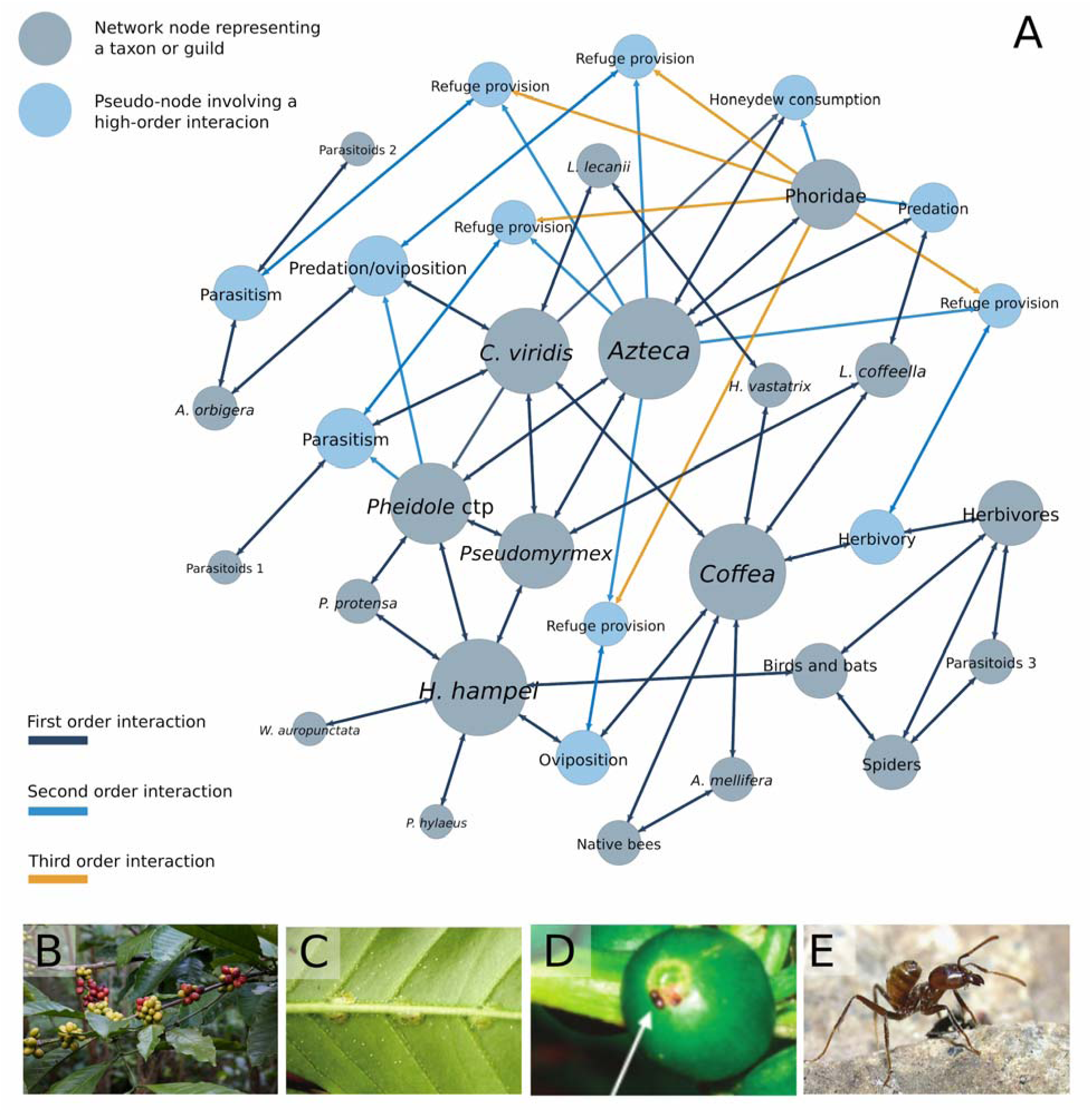
A: Community network with first, second and third order interactions. Grey nodes represent biological taxa and blue nodes are pseudo-nodes representing ecological interactions which are subject to being modified by a HOI. Node size is determined by its degree. First order edges are grey, second order edges are blue and third order edges are orange. B: Coffee plants (*Coffea*). C: Coffee green scale (*Coccus viridis*), a potential pest in the system. D: Coffee berry borer (*Hypothenemus hampei*), one of the main coffee pests, about to penetrate a coffee grain. E: *Azteca* ant, an important regulator of this interaction network. Photographs: Wikimedia Commons by *Jmhullnot* at https://commons.wikimedia.org/wiki/File:CoffeeBerry.jpg (B), John Vandermeer (C, D), Alex Wild (E)

The system here described exhibits different kinds of direct interactions like herbivory and parasitism, but also numerous HOIs (Table S1). For example, *Azteca* ants exert a second order interaction when they inhibit the predation interaction among *C. viridis* and *A. orbygera* by harrasing the latter, mostly without harming it (Vandermeer & Perfecto 2006; Liere & Larsen 2010; Vandermeer et al. 2010). An example of a third order interaction is the effect of the phorid flies, which by paralyzing or chasing away *Azteca* ants, inhibit the second order interaction they exerted and thus enable the predation of *C. viridis* by *A. orbygera* (Hsieh et al. 2012).

### 2.2 Network inference

We used a network approach to analyze the community under study. Species were represented as nodes whose connections were defined by the ecological interactions among them. In order to define the network’s structure, we reviewed published information on this particular agroecosystem and integrated it in a common database.

The reviewing process began with a book that collects over 20 years of research in the area (Perfecto & Vandermeer 2015). All referenced papers that explained, observationally or experimentally, at least one ecological interaction among a pair of species, were examined too. The type of interactions and the direction of their effects were extracted, including qualitative information about their strength, whenever available. If any of the papers in this first group made reference to other investigations in the area, those were also revised. All the information was integrated in a database organized as follows: *transmitter node (*e.g. *H. hampei), recipient node (*e.g. *Coffea*)*, kind of interaction* (e.g. +/-), *description* (e.g. females of *H. hampei* bore into the coffee berries to oviposit and their larvae feed from it) and *reference* (listing of the articles that support the interaction). For HOIs, instead of a *recipient node*, a column was added with the *recipient interaction* (e.g. the presence of *Azteca* prevents *H. hampei* from boring into the coffee, inhibiting herbivory). Interactions that were uncertain, but suspected, were annotated but not considered for the construction of the network. Finally, the network was compared with smaller versions published previously and revised by experts.

### 2.3 Structure definition and general metrics

The structure of the network was visualized with the software *Gephi 0.9.2*. Because network-related methods only contemplate ensembles of nodes connected directly through edges (that is, first order interactions), it is not possible to define a network with edges connecting to other edges, which is the case of HOIs. For this reason, two versions of the network were created: the first one only captured the nodes and their first order interactions; the second one included HOI modified interactions as artificial pseudo-nodes, an artifact that allowed us to use the full force of network theory to analyze the system. Topological analyses were conducted on both versions of the network in order to quantify the effect of HOIs.

The transformation process of HOIs into pseudo-nodes is depicted in Figure 1. Basically, an edge that was affected by a third node was labeled with a new pseudo-node (e. g. a pseudo-node named ‘predation”), so the third node had now a simple edge connecting it to the new pseudo-node. The same logic works for second, third or any higher order interactions. A similar procedure was suggested by Newman (2018), where interactions involving more than two nodes are introduced by adding new nodes belonging to a different category as part of a bipartite graph. This new node is connected by a single edge to each original node. However, this procedure is limited as bipartite graphs do not account for edges between nodes belonging to the same category.

**Figure 1.**
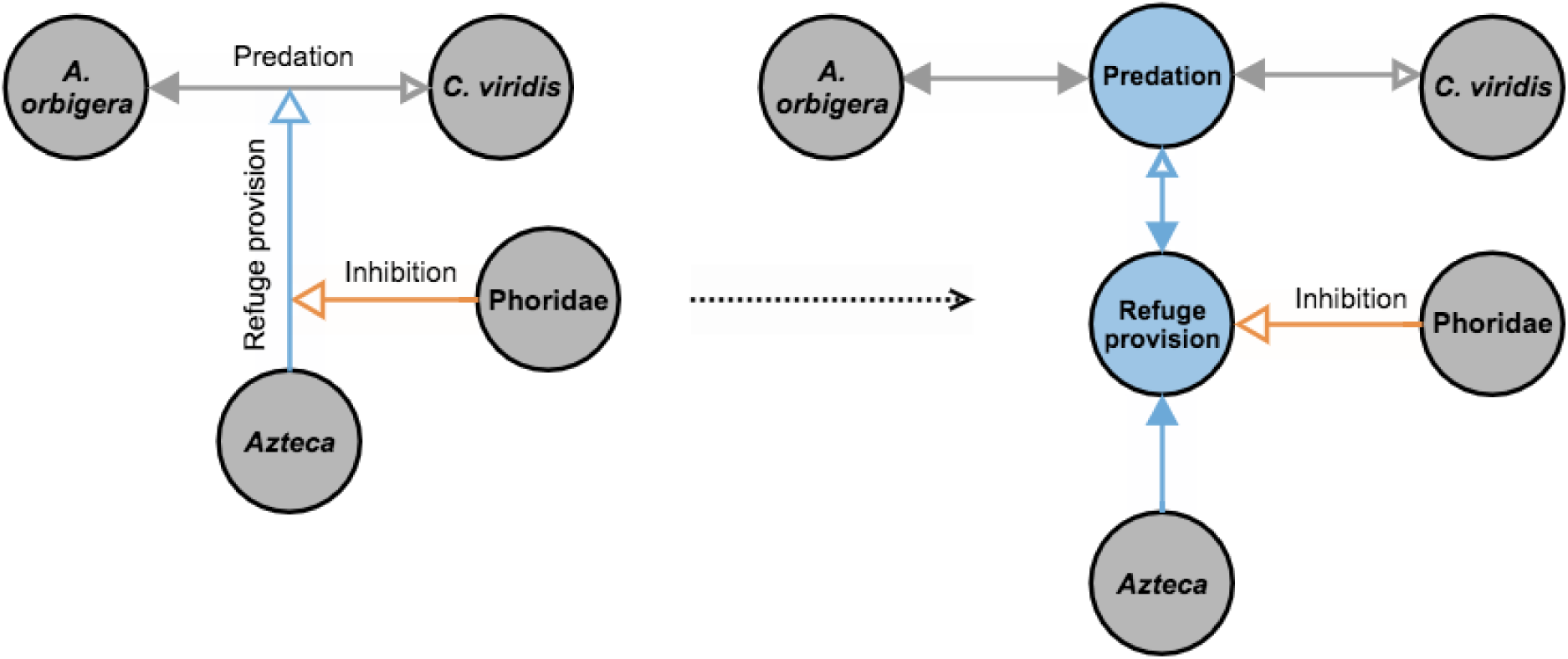
Transformation process of second and third order interactions into pseudo-nodes. The grey nodes represent biological taxa and the blue nodes are pseudo-nodes representing ecological interactions which are modified by a HOI. First order edges are dark grey, second order edges are blue and third order edges are orange. Filled arrows represent positive effects, and unfilled ones, negative effects. For example, the predatory interaction between *A. orbigera* and *C. viridis* is turned into a pseudo-node named *predation* in order to be modified by the refuge provision performed by *Azteca,* a second order interaction. Likewise, this refuge provision is inhibited by the presence of phorids, so it is turned into a second pseudo-node in order to be modified by the third order interaction performed by *Phoridae*.

Once both versions on the network were obtained, standard network metrics were quantified to characterize them: number of nodes, number of edges, mean degree, diameter, density, modularity (using the Louvain algorithm), clustering coefficient, mean path length, and *sigma* and *omega* small world coefficients (Humphries & Gurney 2008; Telesford et al. 2011). Afterwards, we analyzed node relevance according to their centrality. For this, we used four commonly used metrics: degree, closeness centrality, betweenness centrality, and eigenvector centrality. All calculations were made with the software *Gephi 0.9.2*.

### 2.4 The effect of high order interactions on network robustness

We conducted a robustness analysis for both versions of the network (with and without HOIs). Robustness was measured by calculating the area under the curve that depicts the size of the biggest connected component as nodes are removed one by one from the network (Kasthurirathna et al., 2013; Piraveenan et al., 2013; Navarro Díaz 2015). This measure is compared with the area under the curve traced by a complete graph, that is, a graph where every possible pair of nodes is connected by an edge. Thus, following Equation 1, the relationship between these two areas gives us a measure of robustness (for a full derivation of the equation see Piraveenan et al. (2013)).

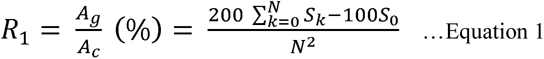

Where A_g_ is the area under the curve of the evaluated graph and A_c_ that of the fully connected graph. *S_k_* is the size of the largest component after *k* nodes have been removed, *S_0_* denotes the initial largest component size, and *N* is the network size. According to the above equation, for a fully connected network of any size, the robustness coefficient (*R*) is always of 100% (taken from Kasthurirathna et al., 2013).

For the empirical network that includes HOIs, only real nodes could be selected for removal, in order to avoid the biologically meaningless action of removing pseudo-nodes. Following this logic, whenever a node got selected for removal, any pseudo-node connected to it was also eliminated, since pseudo-nodes lose their meaning once the species causing the higher order effect is eliminated. Because this modification often resulted in the elimination of several nodes at the time, we modified equation 1 in order to control for it. In the Piraveenan et al (2013) derivation, the area under the curve of the fully connected graph assumes one node removal per step in the *x* axis. If we assume n node removal per step (in order to control for pseudo-node removal in the evaluated graph), this area is *Ac* = *N*2/2*n* and the robustness equation becomes:

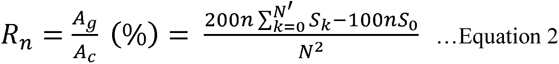

Where *n* is the average number of nodes removed at each step (1.54 in this network) and *N’* is the number of real nodes in the network (*N* minus the number of pseudo-nodes). Equation 1 is equivalent to Equation 2 when n=1 and there are no pseudo-nodes.

Hence, we used equation 1 for the network without HOIs and equation 2 for the network with HOIs. For each of these networks, two node removal methods were tested. With the first one, nodes were randomly selected and removed one by one until removing them all. This was done 200 independent times and a robustness average was obtained. The second method consisted of removing nodes by degree, from highest to lowest.

The robustness of each network with each removal method was also compared with the average robustness of 200 randomized but comparable networks, i.e. with the same number of nodes, average degree or interaction density. Three types of random networks were used: totally random networks (Erdös & Rényi 1960), small-world networks (Watts & Strogatz 1998) and scale-free networks (Barabási & Albert 1999). The first model generates random networks from a set of nodes in which the edges are independently created between any pair of nodes with a probability *p*. Because the structure of ecological networks is far from being random, we also used small-world and scale-free networks, which have been proved to share structural characteristics with many real world networks (Montoya & Solé 2002; Barabási & Bonabeau 2003). Small-world networks follow an algorithm that starts with a regular lattice where each node is connected to its *k* closest neighbors, and where each edge is then re-connected to a randomly chosen node with a certain probability, avoiding duplicates and self-loops. This construction produces networks with a high clustering coefficient and short paths, two particularities that have been found in many ecological webs (Montoya & Solé 2002). The last method builds networks with a preferential attachment mechanism, where nodes are added sequentially such that each new node is connected to a number *m* of existing nodes, where the probability to choose a node for connection is proportional to the number of links that this node already has. This creates networks with power-law degree distributions, another characteristic that has been widely found in ecological webs (Barabási & Bonabeau 2003). For the Erdös Rényi method we used the values *N*=34 and *p*=0.095, and *N*=22 and *p*=0.145 for networks representing cases with and without HOIs, respectively (where *N* is the number of nodes of the empirical web and *p* is taken from their density). For the Watts-Strogatz method, we chose *N*=34, *k*=3 and *p*=0.5, and *N*=22, *k*=3 and *p*=0.5 for networks representing cases with and without HOIs, respectively (where *k* is the average degree of the empirical web and *p* was arbitrarily chosen). For the Barabasi-Albert method we chose *N*=34 and *m*=1, and *N*=22 and *m*=2 for networks representing cases with and without HOIs, respectively (where *m* is chosen so that the resulting average degree matches the empirical average degree).

Because nodes in the empirical network with HOIs were removed along with their associated pseudo-nodes as discussed above, the randomized versions of this network needed to emulate this process too. This was done in the following way: First, we quantified the probability to remove a number *n* of pseudo-nodes with each real node removal in 100 simulations of the empirical network with HOIs. Then, in the randomized networks (composed of 34 nodes), a subset of 22 randomly chosen nodes was defined to stand for the real nodes, while the remaining 12 nodes stood for the pseudo-nodes. At each removal step, a node was removed (randomly or by degree as explained above) from the real nodes pool alongside with *n* nodes from the pseudo-node pool, with *n* drawn from the probability distribution derived from the mentioned simulations. Again, we used Equation 1 for calculating robustness of the randomized versions of the network without HOIs and Equation 2 for the randomized versions of the network with HOIs. With these numerical experiments we were able to compare, on the one hand, the robustness of the two versions of our network, that is, with and without HOIs, and on the other hand, each empirical robustness with their randomized analogues. Simulations were done with the library NetworkX 2.5 (Hagberg et al. 2008) in *Python 3.7.1*. and ANOVA tests were performed in *RStudio 1.2.1335* (RStudio Team 2020). Scripts are publicly available at: https://github.com/laparcela/CoffeeNetworkStructure

## 3. Results

### 3.1 Network inference

From literature revision, 48 interactions between 22 nodes were established out of 44 scientific papers and books, all conducted in our study site (Figure 2). This information is organized in the supplementary material table S1.

**Figure 2.**
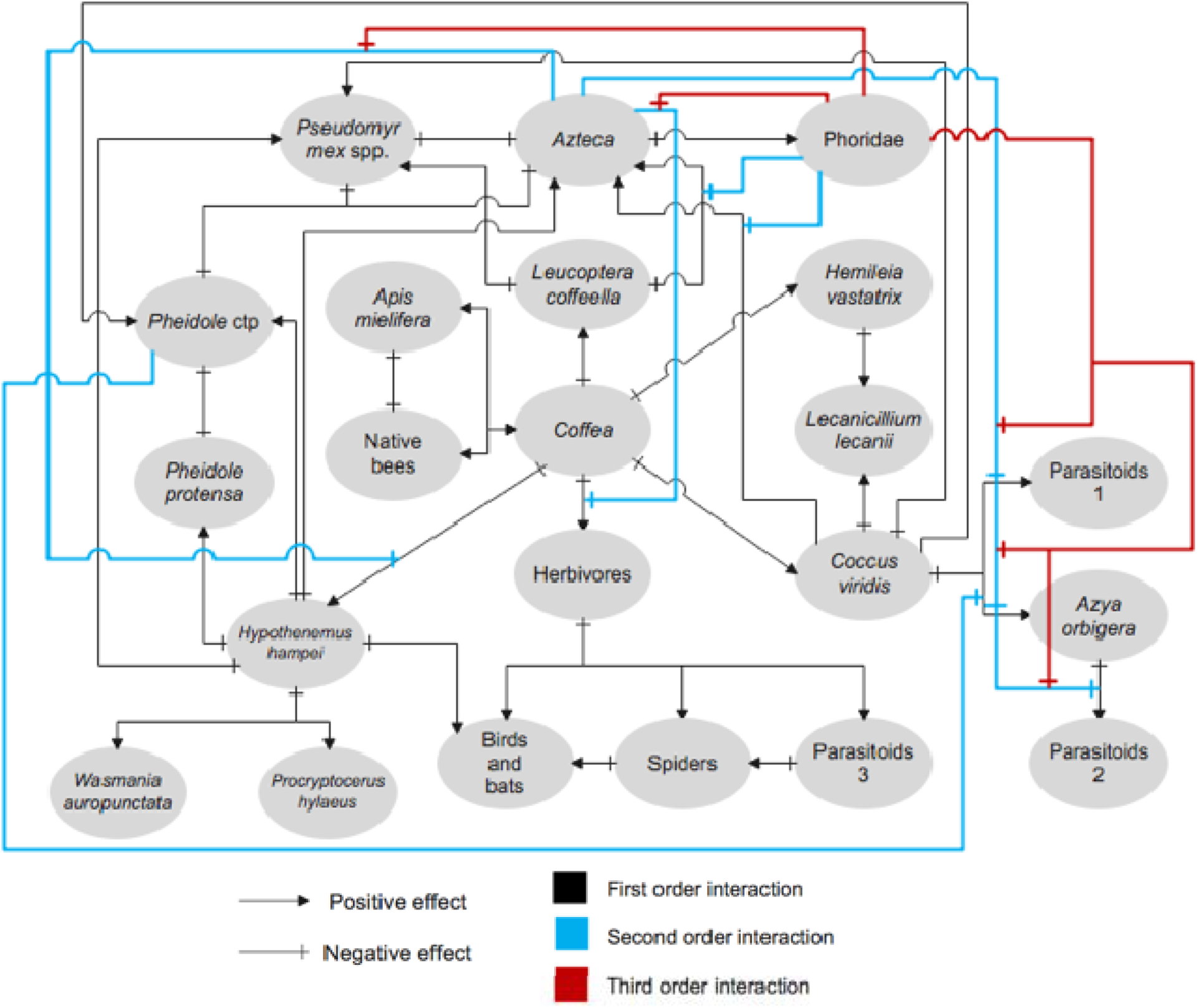
Complete network before transformation from HOIs to pseudo-nodes. Black lines are first order interactions, blue lines are second order interactions and red lines are third order interactions.

### 3.2 Structure definition and general metrics

Two versions of the web were obtained with *Gephi,* the first one containing only first order interactions and the second one after adding pseudo-nodes for HOIs (Figure 3).

Without HOIs, the network is composed of 22 nodes and 68 interactions, while incorporating HOIs makes it a network of 34 nodes and 104 interactions. Both networks have an approximate average degree of 3. Table S2 summarizes the general metrics obtained for both versions of the network. Centrality analysis showed that *C. viridis, Coffea, H. hampei, Azteca, Pheidole* ctp. and *Pseudomyrmex* spp are the nodes with the highest rankings in both networks and for different centrality metrics (Figure 4).

**Figure 4.**
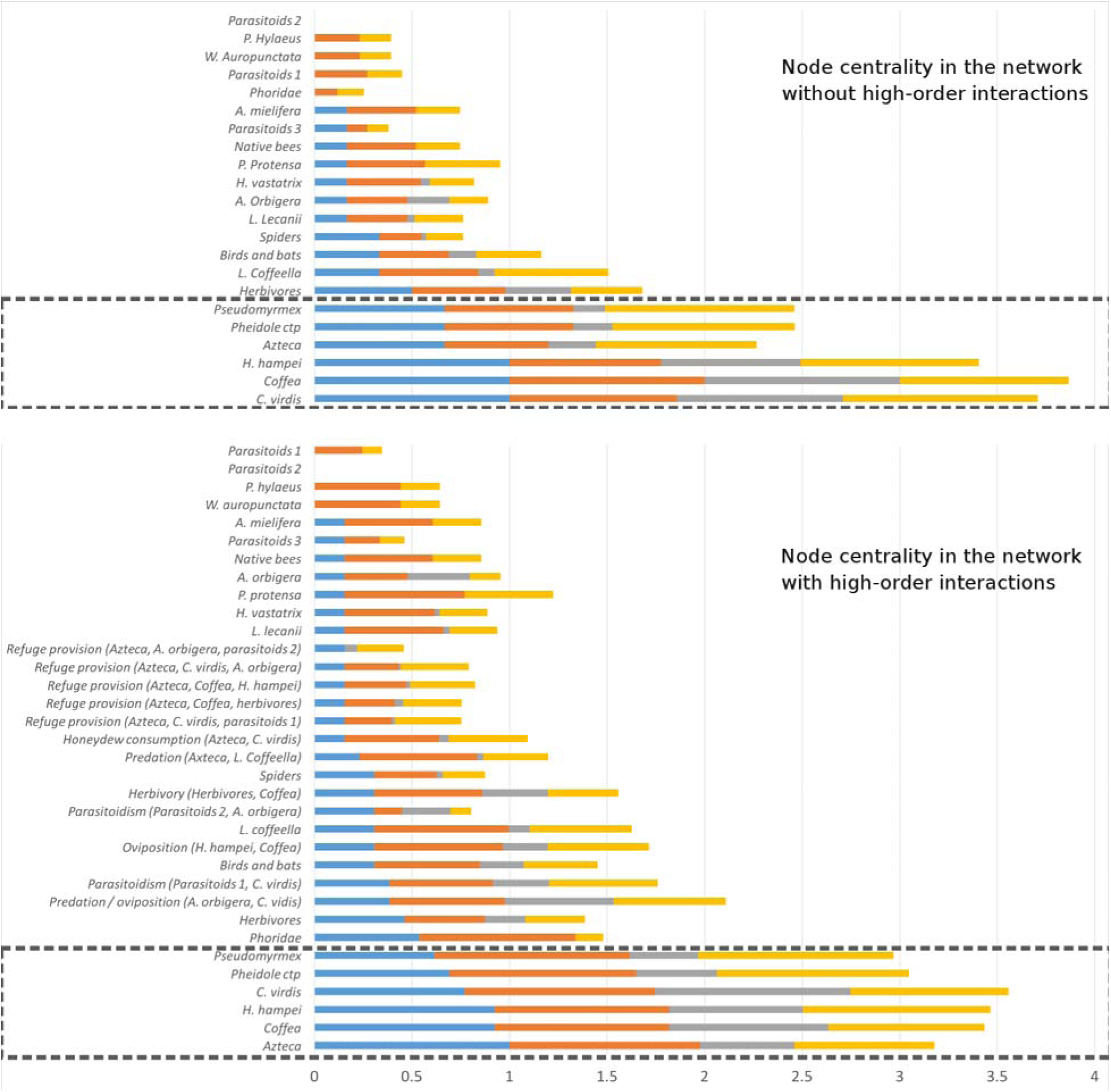
Node centrality analysis for the network without HOIs (above) and with HOIs (below). *C. viridis, Coffea, H. hampei, Azteca, Pheidole* ctp and *Pseudomyrmex* spp are the highest ranking nodes in both networks.

### 3.3 The effect of high order interactions on network robustness

Figure 5 presents the results of the robustness analyses for the empirical coffee networks with and without HOIs, as well as the results for the three different types of randomized networks with comparable structures. In the case of the empirical networks, the addition of HOIs did not significantly change the network robustness under random node removal, but robustness increased significantly under directed node removal. In contrast, for the three types of randomized networks subject to the two node removal protocols, networks with comparable structures to those with HOI addition significantly lost robustness, except for the completely random networks (Erdos-Renyi) under random removal, which showed no significant changes. Additionally, in the node removal by degree, taking HOIs into account made the empirical network more robust than all its randomized counterparts. Statistical analyses can be found in the table S3 of the Supplementary material. Because all randomized analogues of the network with HOIs have the tendency to lose robustness, while the robustness of the actual empirical networks is either unchanged or increased by HOIs, we can say that the effects observed in the empirical networks are indeed a result of HOI addition and not of simply increasing the number of interactions. Indeed, it seems that high order interactions favor robust network structures that may enable the coexistence of diverse systems.

**Figure 5.**
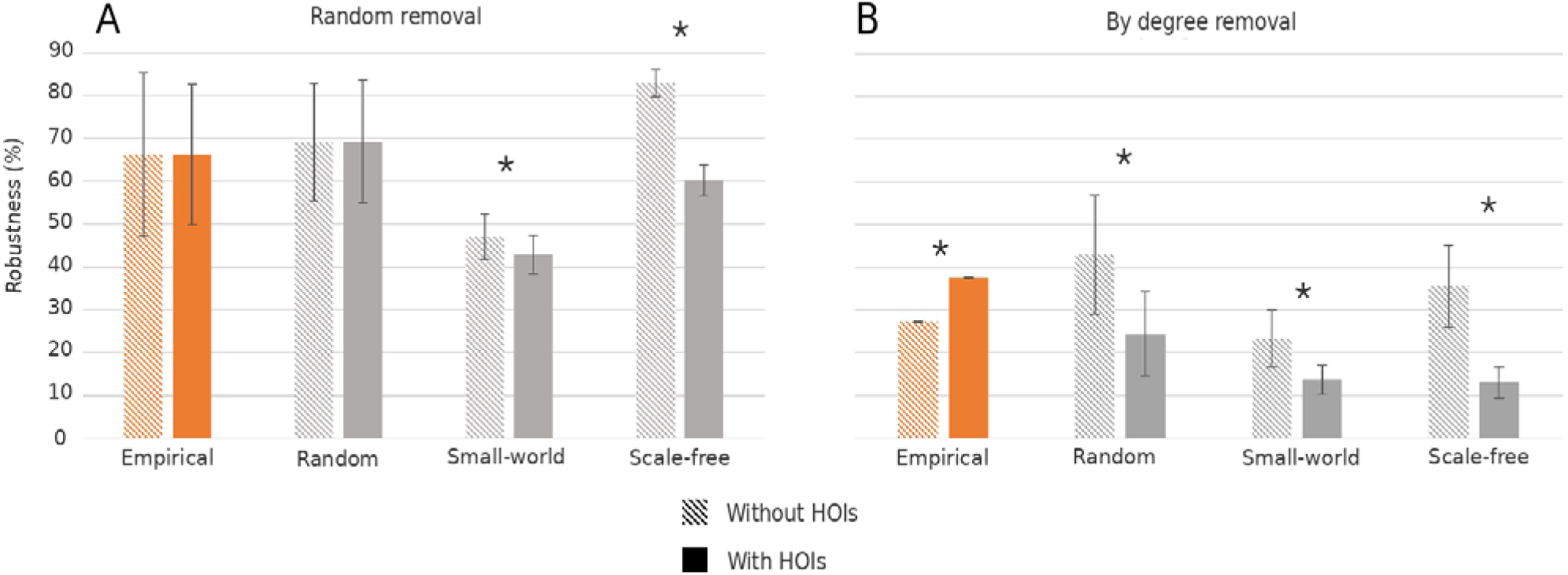
Robustness of the coffee-associated network, with and without HOIs, as well as random, small-world and scale free networks with same *n*, mean degree and density. A: Under random node removal, the empirical web (orange bars) and the totally random networks (grey, left) are not significantly changed by the addition of HOIs; while small-world (grey, middle) and scale-free networks (grey, right) loose robustness under HOI addition. B: When removing nodes by degree, the empirical network (orange bars) is significantly more robust when HOIs are added, while the three types of randomized networks (grey bars) lose robustness when their structures are comparable to that with HOI addition.

## 4. Discussion

We have integrated a vast set of empirical evidence into a coffee-associated network that includes both simple and high order ecological interactions (Figure 3). This network has enabled us to test the role of HOIs on the network’s structural robustness for a system of great ecological and agricultural importance. We find that the robustness of the coffee-associated network structure is unchanged or increased by HOI addition, and random reconfigurations indicate that this effect is not simply due to edge addition (Figure 5). This goes in agreement with previous studies considering hypothetical networks and different measures of system function like stability or feasibility, where the addition of simple interactions has been found to negatively repercute on system function while HOI addition has a neutral or a positive effect (May 1972; Bairey et al., 2016; Grilli et al., 2017; Singh & Baruah, 2020; Li et al., 2020). Our results therefore support the idea that HOIs contribute to the maintenance of highly diverse ecological communities.

In our study, the structural robustness of the network was evaluated with the change in size of the biggest connected component as the nodes were gradually removed, at random or by targeting nodes of higher degree first. This way of conceptualizing robustness assumes that the connection between network components is related to the function and integrity of the system, implying that a fully connected network can maintain its elements and overall functions better than a disaggregated or partially disconnected network (Albert et al. 2000; Dekker & Colbert 2004; Piraveenan et al. 2013; Sheykhali et al. 2020). Indeed, previous work on the coffee agroecosystem for which the network under study has been uncovered suggests that some agroecosystemic functions, such as pest control, rely on the dynamics of the whole system and on the documented interactions taking place (Vandermeer et al. 2010). In the particular case of agroecosystems, the integrity of the network, in other words the maintenance of its diversity, is also likely to be associated to yield and yield stability in the face of diverse perturbations (Gaudin et al. 2015; Manns & Martin 2018).

While the coffee-associated system was studied as an undirected network, the type and sign of its HOIs could inform the mechanism through which HOIs affect the overall robustness. In ecological terms, the HOIs considered in this network may act as buffers of the interactions they modify, thereby diminishing their intensity. For example, refuge provisioning, where one species protects another from one or several predators, may not only explain prey survival (which is important for maintaining the predator), but also how predators avoid competitive exclusion. It is possible that these mechanisms, coupled with spatial and temporal heterogeneity, may create the necessary conditions for coexistence. However, it is important to bear in mind that individual HOIs may have effects in different directions. Specially in the case of agroecosystems, where effects are measured also in terms of human-based values like productivity, the effect of individual HOIs should not be universally assumed as positive. For instance, it has been shown that the ant *Wasmannia auropunctata* can indirectly protect the coffee leaf miner against potential predators, potentially limiting the effectiveness of biological control elements (Perfecto et al., 2021).

The structural analyses of the coffee-associated network also allowed us to identify nodes with high centrality according to different metrics (Figure 4). Centrality has been used as an indicator of the role of individual nodes in the overall dynamics of networks; in ecological networks, node centrality is thought to reflect how a node contributes to the flow of energy and matter and ecosystem functioning. Highly central nodes indirectly connect many other nodes in a network and act as ‘bridges’, a reason why centrality has been amply used in the study of socio-ecosystems networks (Freeman, 1979; Raghavan Unnithan et al., 2014; Lü et al., 2016; Horcea-Milcu et al., 2020; Arroyo-Lambaer et al., 2020). We identified five nodes that systematically exhibited a high centrality, independently of the centrality measure: *C. viridis, Coffea, H. hampei, Azteca, Pheidole* ctp. and *Pseudomyrmex* spp. This is in agreement with the crucial role of the coffee plant in this agroecosystem, as well as the effect of its potential pests and pest enemies in its growth and development (Vandermeer et al. 2010). However, at this point we cannot rule out the possibility that the high centrality of these nodes is due to a bias in sampling and research efforts. We are currently pursuing analyses that go beyond the study of the network structure and that might help uncover the role of HOIs and highly central nodes on the dynamics of populations on such a network.

To conclude, our results support the hypothesis that HOIs can contribute to the maintenance and robustness of highly diverse ecological systems, and agroecological systems in particular. In agreement with previous empirical and theoretical studies, our work points to the importance of agroecological management and practices that are based on a deep ecological understanding of productive systems, as well as to the importance of a high diversity of taxons and interactions for the robustness and functioning of agroecosystems.

## Acknowledgements

The authors thank Blanca Hernández Hernández and Yolanda Robledo Arratia for formatting and figure edition. We also thank members of *La Parcela* Laboratory and the two reviewers for their valuable comments and suggestions. This article covers part of the requirements to obtain the PhD Degree in the Posgrado en Ciencias Biológicas de la Universidad Nacional Autónoma de México.

## 8. Supplementary Material

**Table S1.**
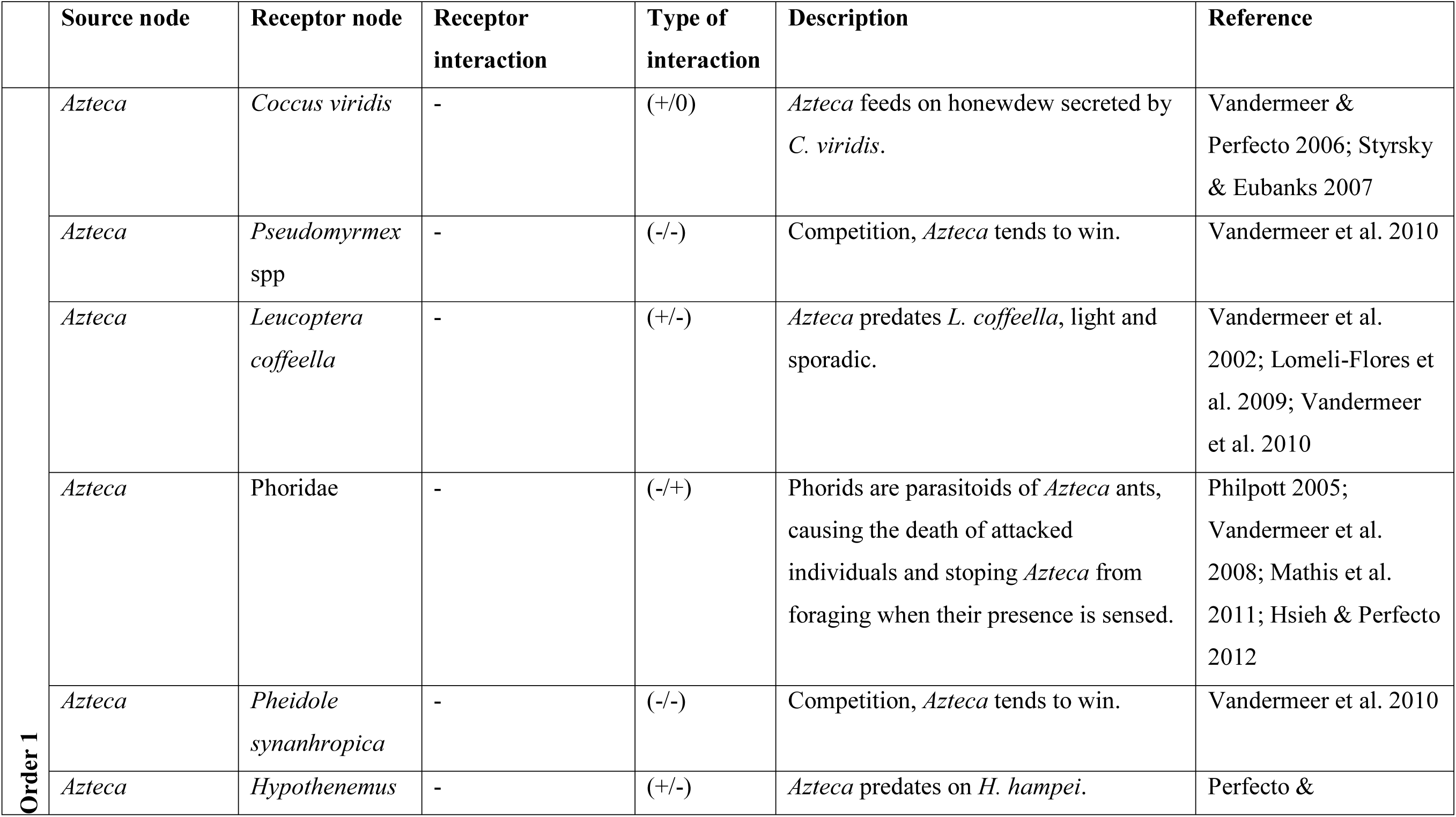
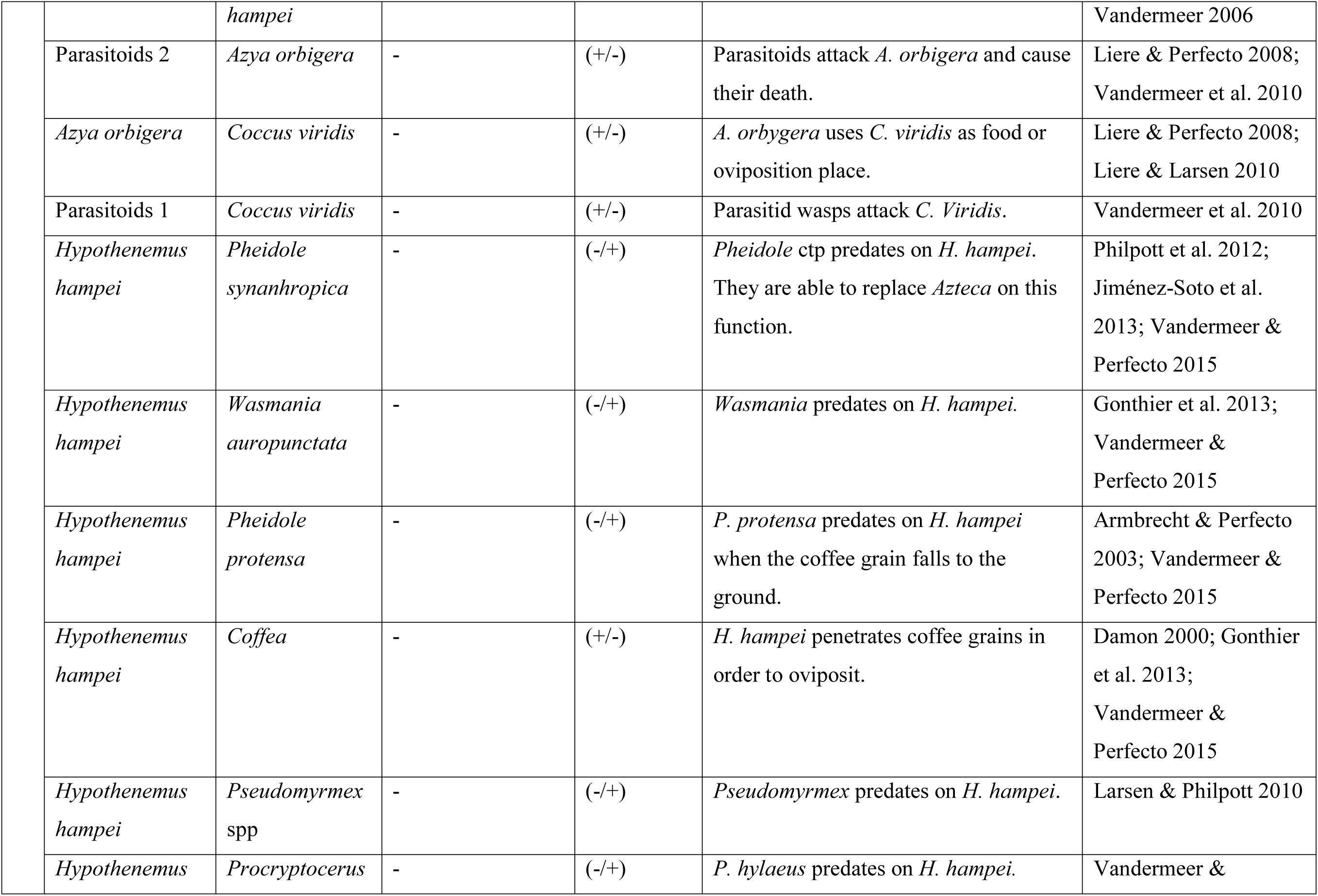
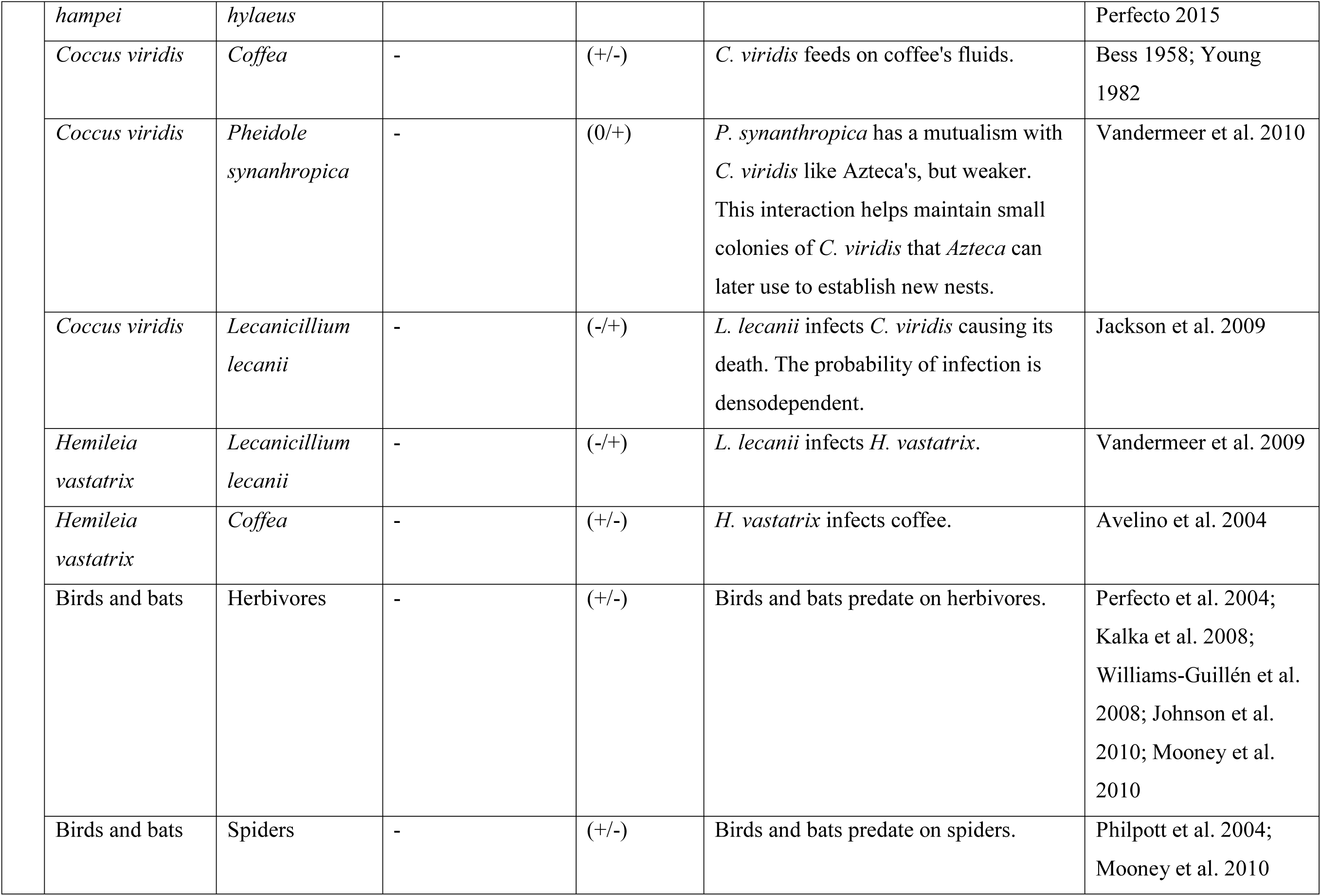
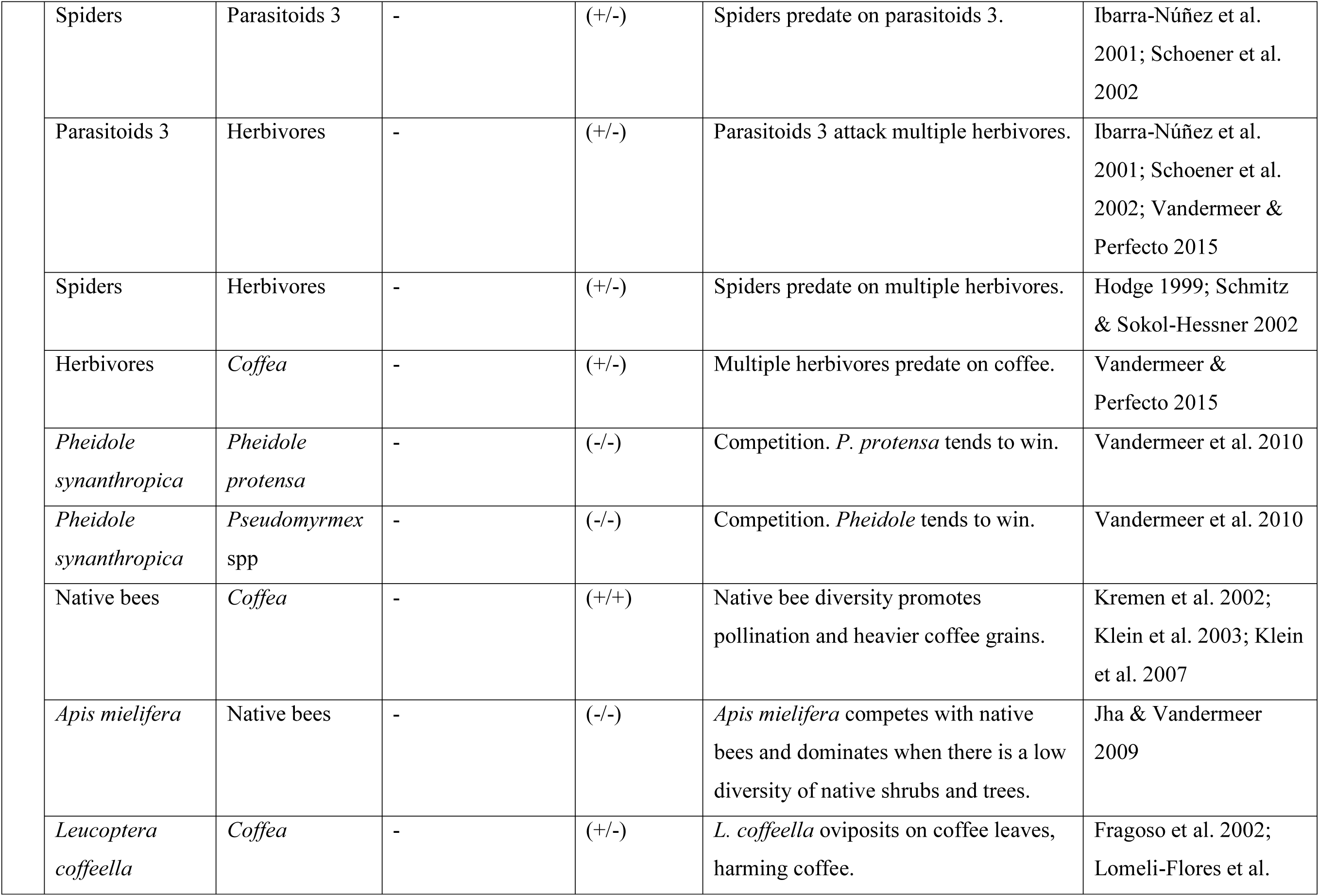
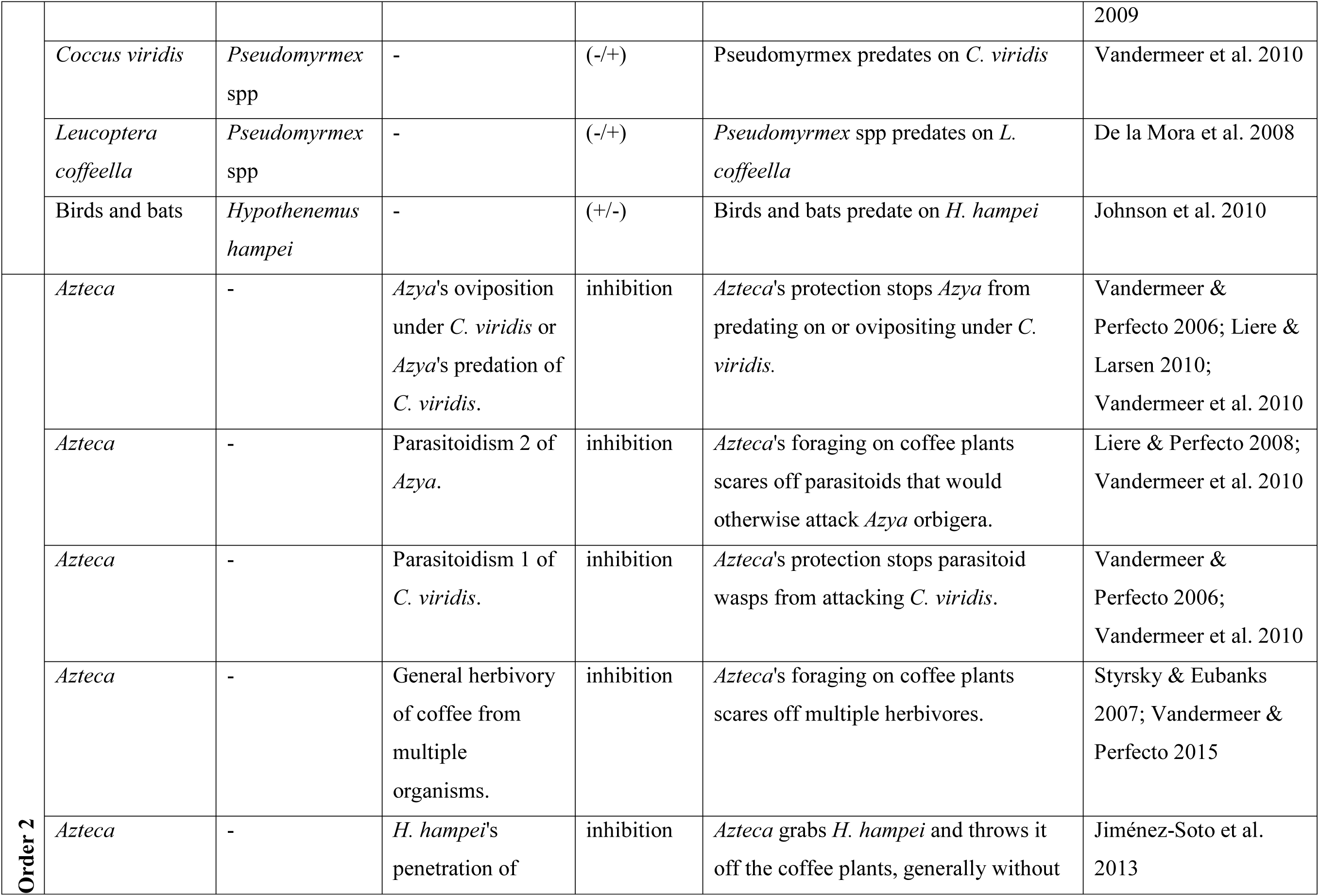
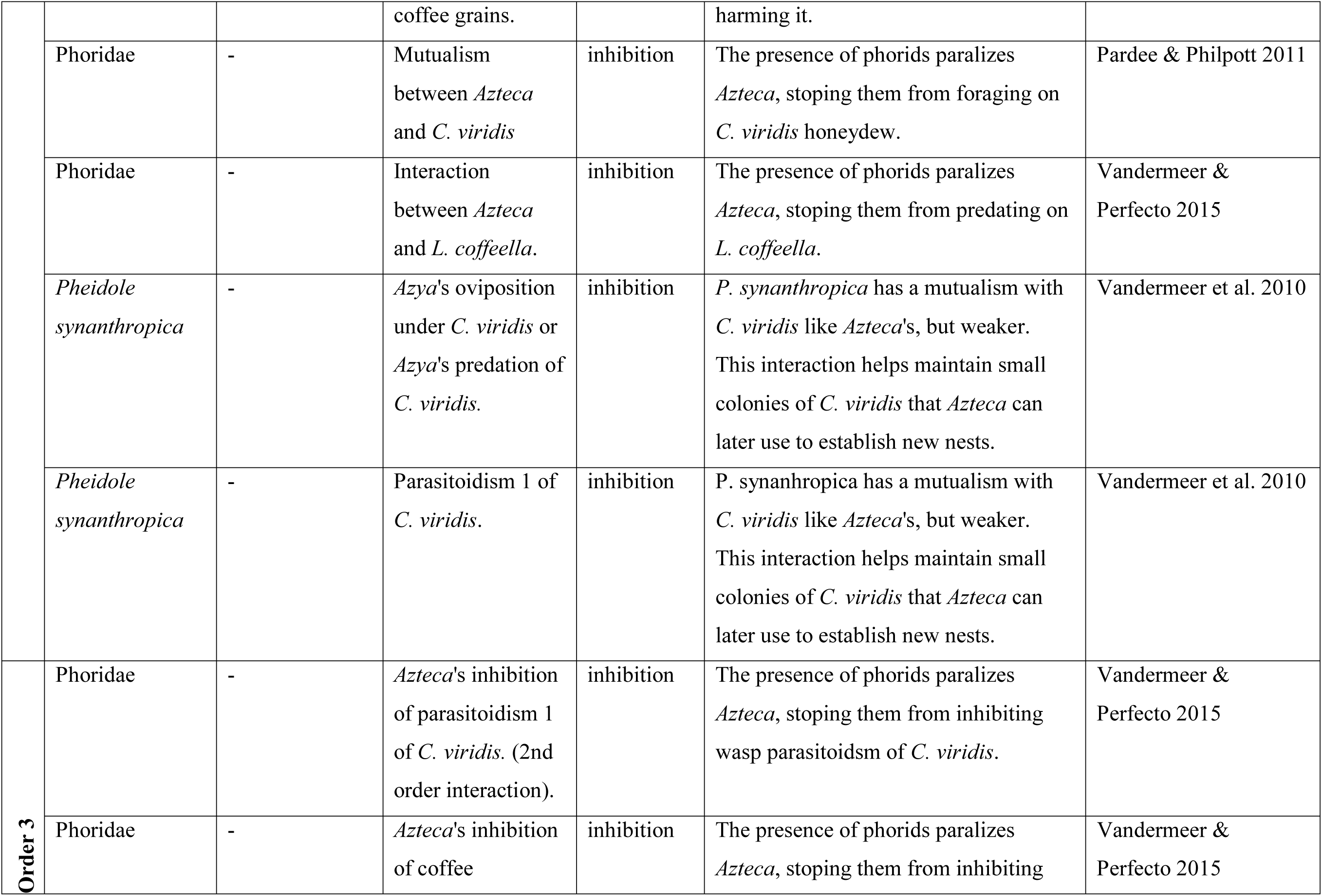
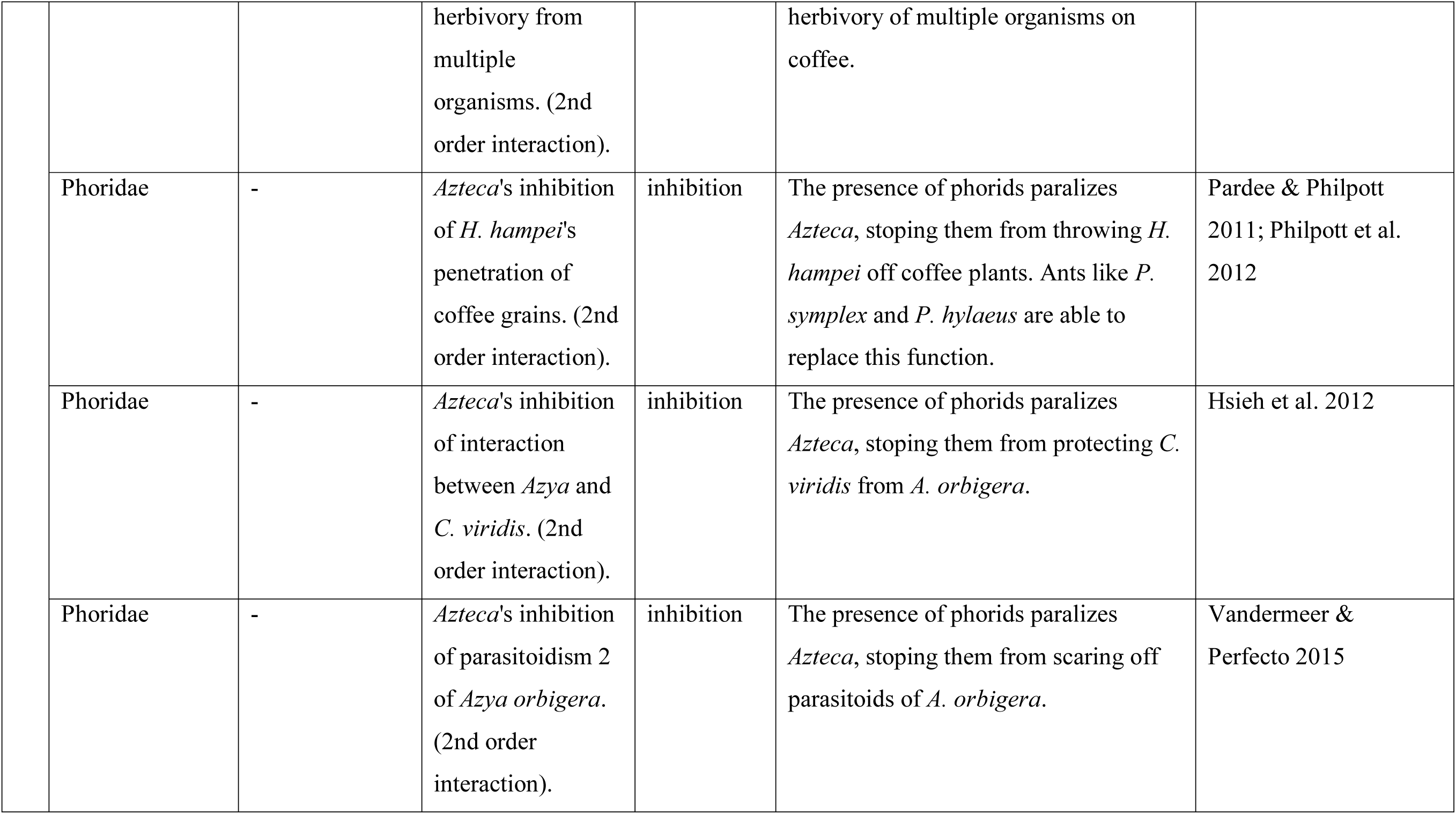
Complete database containing all nodes and interactions included in the network, as well as references supporting them.

**Table S2.** General metrics obtained for the network with and without HOIs.

|  | *Without HOIs* | *With HOIs* |
| --- | --- | --- |
| Nodes | 22 | 34 |
| Edges | 68 | 104 |
| Average degree | 3.091 | 3.059 |
| Diameter | 5 | 8 |
| Density | 0.147 | 0.093 |
| Modularity | 0.44 (5 modules) | 0.45 (6 modules) |
| Clustering coefficient | 0.334 | 0.261 |
| Average path length | 2.632 | 3.362 |
| Small world coefficient (*sigma*) | 1.36 | 1.4 |
| Small world coefficient (*omega*) | 0.33 | 0.51 |

**Table S3.** ANOVA analyses for the robustness of all networks.

|  |  | Degrees of freedom | Sum of squares | Mean sum of squares | Statistic | *p* value | *eta* square | *omega* square | *epsilon* square |
| --- | --- | --- | --- | --- | --- | --- | --- | --- | --- |
| Random node removal | Empirical with HOIs vs without HOIs | 1 | 0.019 | 0.019 | 0 | 0.994 | 0 | -0.003 | 0 |
|  | Random with HOIS vs without HOIS | 1 | 0.014 | 0.014 | 0.17 | 0.68 | 0 | -0.002 | -0.002 |
|  | Small-world with HOIS vs without HOIS | 1 | 1774.66 | 1774.66 | 72.14 | <0.001 | 0.153 | 0.151 | 0.426 |
|  | Scale-free with HOIS vs without HOIS | 1 | 50986.63 | 50986.63 | 4463.04 | <0.001 | 0.918 | 0.918 | 3.349 |
| By degree node removal | Random with HOIS vs without HOIS | 1 | 34281.86 | 34281.86 | 236.54 | <0.001 | 0.373 | 0.371 | 0.771 |
|  | Small-world with HOIS vs without HOIS | 1 | 9312.47 | 9312.47 | 333.91 | <0.001 | 0.456 | 0.454 | 0.916 |
|  | Scale-free<br>with HOIs<br>vs without<br>HOIs | 1 | 50922.49 | 50922.49 | 964.08 | <0.001 | 0.708 | 0.707 | 21022.325 |

